# AptaBERT: Predicting aptamer binding interactions

**DOI:** 10.1101/2023.11.24.568626

**Authors:** Flemming Morsch, Iswarya Lalitha Umasankar, Lys Sanz Moreta, Paridhi Latawa, Danny B. Lange, Jesper Wengel, Huram Konjen, Christian Code

**Affiliations:** Dianox, Fruebjergvej 3, 2100 Copenhagen, Denmark; Technical University of Denmark, Anker Engelunds Vej 101, 2800 Kongens Lyngby, Denmark; Massachusetts Institute of Technology, 77 Massachusetts Avenue, Cambridge, MA 02122, United States; Google, 1600 Amphitheatre Parkway, Mountain View, CA 94043, United States; University of Copenhagen, Noerregade 10 & Blegdamsvej 3B, 1165 & 2200, Copenhagen, Denmark

**Keywords:** aptamer, nucleic acid, DNA, RNA, oligonucleotides, Large Language Model (LLM), Machine Learning, Deep Learning, protein interaction, drug discovery

## Abstract

Aptamers, short single-stranded DNA or RNA, are promising as future diagnostic and therapeutic agents. Traditional selection methods, such as the Systemic Evolution of Ligands by Exponential Enrichment (SELEX), are not without limitations being both resource-intensive and prone to biases in library construction and the selection phase. Leveraging Dianox’s extensive aptamer database, we introduce a novel computational approach, AptaBERT, built upon the BERT architecture. This method utilizes self-supervised pre-training on vast amounts of data, followed by supervised fine-tuning to enhance the prediction of aptamer interactions with proteins and small molecules. AptaBERT is fine-tuned for binary classification tasks, distinguishing between positive and negative interactions with proteins and small molecules. AptaBERT achieves a ROC-AUC of 96% for protein interactions, surpassing existing models by at least 15%. For small molecule interactions, AptaBERT attains an ROC-AUC of 85%. Our findings demonstrate AptaBERT’s superior predictive capability and its potential to identify novel aptamers binding to targets.

## 1 Introduction

### 1.1 Use and selection of aptamers

Aptamers, consisting of short single-stranded DNA or RNA sequences, have garnered significant attention for their ability to bind to a diverse array of targets, from small molecules [1] to proteins [2], viruses and cells [3, 4]. They offer specificity and affinity comparable to antibodies [5, 6], positioning them as ideal candidates for many applications in diagnostics, research tools, and therapeutics. Emerging as alternatives to antibodies, aptamers facilitate antibody-free detection of pathogens and toxins [7, 8] and serve as valuable molecular probes. Research indicates that combining multiple aptamers can augment binding efficacy and specificity [9]. Unlike antibodies, aptamers possess a structure-switching ability, which has been harnessed for early cancer diagnostics [10, 11] and sickle cell anemia detection [12], as well as detecting environmental and food contaminants [13]. Lastly, the FDA has thus far approved two therapeutic aptamers: Macugen and Izervay [14, 15].

Aptamer research began in the 1990s and the Systemic Evolution of Ligands by Exponential Enrichment (SELEX) process has been central to selecting aptamers specific to targets. The method starts with a library of DNA or RNA sequences, including a random section (N10-N60 nucleotides) flanked by primer regions. Specific aptamers that bind to the target are enriched and often truncated post-selection to enhance stability and binding. Binding and specificity assays follow to determine the dissociation constant and specificity [16]. To improve SELEX efficiency, various techniques have emerged, such as Capillary Electrophoresis SELEX, Cell SELEX, and Microfluidic SELEX [17].

However, SELEX has drawbacks, including library and selection biases, which may hinder identifying the optimal aptamer. Libraries tend to have a nucleotide bias towards guanine-rich sequences [18] and against adenosine inclusion [19]. During SELEX, certain aptamers may be preferentially selected due to PCR or amplification biases, potentially overlooking better binders. The laborious nature of measuring dissociation constants compounds the issue if many aptamers are chosen [20]. Notably, the guanine-rich aptamer AS1411 which binds to nucleolin and is well characterized as a potential therapeutic for cancer was discovered serendipitously [21], not through SELEX, highlighting the process’s limitations despite three decades of refinement [22].

Computational advances, including bioinformatics and machine learning, have improved the characterization of aptamers and their targets post-SELEX using tools like Aptasuite [23], RNAComposer [24], and 3DNA [25]. Nevertheless, these state-of-the-art tools have yet to supplant SELEX.

### 1.2 Data scarcity obstructs the development of reliable algorithms

The scarcity of data hampers the development of reliable algorithms for aptamer research. Machine learning has shown promise in predicting aptamer-protein interactions [26], generating aptamer structures [27], optimizing SELEX protocols [28], and understanding aptamer folding behavior [29]. Yet, the lack of data, with only ∼ 1400 data points in public databases (https://sites.utexas.edu/aptamerdatabase/), limits the full exploitation of modern deep learning architectures that require larger datasets for reliable generalization [30, 31, 32]. This paucity contrasts with the abundance of data for proteins and small molecules [33, 34, 35, 36, 37] and makes building generative models for aptamers an arduous task. To address the data limitation in computational aptamer research, we introduce a novel method that utilizes a BERT-based architecture trained on Dianox’s extensive dataset to predict aptamer-target interactions.

## 2 Related work

Various approaches have attempted to predict aptamer structures and interactions. Physicochemical property-based methods require knowledge of both the aptamer and protein structures to predict their interaction in the three-dimensional space. Aptamers, existing in secondary or tertiary forms, present a highly flexible complex that facilitates high-affinity interactions with a wide range of targets [1, 2, 4]. Chen et al. assessed five algorithms − CentroidFold, Mfold, RNAfold, RNAStructure, and Vfold2D − for predicting aptamer secondary structures, achieving average accuracies above 75% [38]. This suggests partial reliability in predicting secondary structures. Additionally, other software packages like seqfold are also available for this task [39]. Models such as AptaTRACE [40] and APTANI [41] focus on clustering aptamer sequences by secondary structure motifs, which can streamline SELEX experiments by integrating these motifs into their experimental design. This computational angle is becoming increasingly relevant due to the limitations inherent in the SELEX process.

### 2.1 Modeling aptamer-target interactions

Moving to the modeling of aptamer-target interactions, machine learning has been pivotal, with models emphasizing feature engineering through various methods, including pseudo-nucleotide compositions [42, 43, 44], pseudo-amino acid compositions [45, 46, 26], and the position-specific score matrix of proteins [47], supplemented by the physicochemical properties of the aptamers and proteins involved [42, 43, 44, 45, 46, 26, 47, 48, 49, 50, 51, 52]

The architectures employed range from random forests [47, 45, 44] to standard multi-layer perceptrons (MLP) [26, 53], aiming to predict binary outcomes of aptamer-protein interactions. Due to a scarcity of negative interaction data, all the aforementioned studies resorted to using randomized sequences as a control. The binding affinity, measured by the dissociation constant (*K*_*d*_), is crucial for discerning the strength of these interactions, where lower *K*_*d*_ values signify stronger affinities [3, 54]. For machine learning applications, establishing a *K*_*d*_ threshold is crucial to transform continuous affinity measurements into binary classification outcomes.

### 2.2 Rapid improvement in new deep learning architectures enables reliable transfer learning

The surge in large deep learning architectures like transformers, variational autoencoders (VAEs), and diffusion models has revolutionized data processing and learning from extensive datasets [55, 56, 57, 58, 59]. These foundational models, including Large Language Models (LLMs), have significantly impacted our ability to derive meaningful representations from complex datasets [31, 60, 61]. Yet, the application of these models to biological data remains challenging due to its complex and often scarce nature. To date, RaptGen is the only known method employing a VAE architecture to generate aptamers from SELEX data [27]. The scarcity of models capable of leveraging LLMs pre-training inspired the development of a novel method in this project, which is based on the transformer model architecture of BERT [30].

## 3 Dataset

In this study, we sample a subset of Dianox’s database consisting of 67552 aptamer-target pairs, of which 9.67% are labeled sequences. Initially, we utilize the entire database, disregarding the labels, to train the Masked BERT model and to gather the learned sequence pair representations. Subsequently, we employ those learned representations to perform supervised training for aptamer-target interaction prediction using only the labeled data with either protein (*Protein* dataset) or small molecule (*Chemical* dataset) targets. As illustrated in figure 1c the targets predominantly consist of protein structures, and the aptamer sequences are mostly RNA-based (figure 1d).

**Figure 1.**
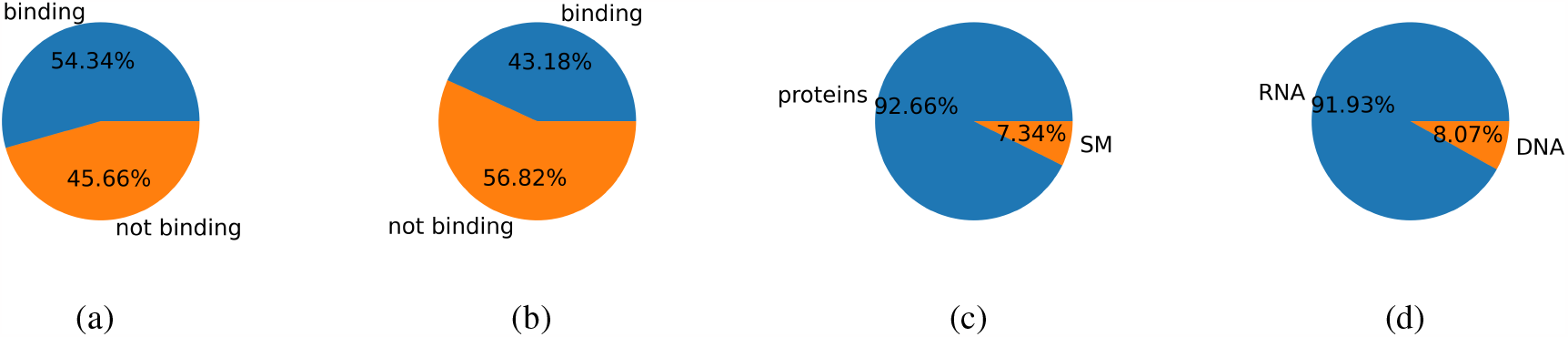
Dataset composition. a) shows the *Protein* dataset distribution of data labels within the labeled subset of the aptamer-protein dataset. Orange represents non-binding aptamer-protein pairs, while blue represents binding pairs. b) depicts the *Chemical* dataset distribution of data labels in the labeled subset of the aptamer-chemical dataset, with orange indicating non-binding pairs and blue indicating binding pairs. c) illustrates the target category distribution, distinguishing between proteins and small molecules (SM). d) displays the distribution of aptamer classes, highlighting that the majority of the aptamers are RNA-based molecules.

The complete dataset comprises i) aptamer sequences with lengths ranging from 6-150 bases and ii) targets with lengths varying from 18-3685 amino acids for proteins and 4-814 SMILES characters for the chemical molecules.

To perform fine-tuning and evaluate the predictive power of the model, we use only the subsets of labeled aptamer-target pairs. The *Chemical* dataset, comprising aptamer-chemical molecule pairs, contains 1577 samples with 43.18% binding sequences. The *Protein* dataset includes 4950 samples, with 54.34% representing binding pairs. The label proportions are illustrated in figures 1a and 1b.

Class assignment for the aptamer-target pairs is based on a dissociation constant *K*_*d*_ threshold set at 10nM. Pairs with a *K*_*d*_ lower than this threshold are categorized as having a positive interaction, indicative of binding, whereas those with a *K*_*d*_ exceeding the threshold are classified as non-binding, reflecting a negative interaction.

## 4 AptaBERT

AptaBERT employs a two-step architecture starting with pre-training an unsupervised Masked LM, followed by fine-tuning with supervised classification on a labeled dataset. This method aims to learn a combined sequence representation that could potentially embed information about molecular interactions between nucleic acids and their targets, encompassing proteins and small molecules. During preprocessing, the model is trained for 100 epochs, amounting to 337500 steps. Subsequently, fine-tuning is conducted using a small neural network on a dataset of aptamer-protein pairs (AptaBERT Prot) and aptamer-small molecules (AptaBERT Chem). Each subset utilizes the same representations learned by the pre-trained BERT MaskedLM model and is trained five times for 15 epochs. The five models are then aggregated into an ensemble model as detailed in section 4.5.

### 4.1 BERT as a foundational model

BERT (Bidirectional Encoder Representations from Transformers) [30] comprises 12 encoding layers, each following the original transformer architecture [55]. Designed to generate embeddings that represent words, which in this study correspond to processed aptamer-target pairs, BERT incorporates attention information within a certain context. Transformer-based models leverage scaled dot-product attention [62] and positional embeddings to process long sequences (long range dependencies preservation) and to generate a weighted average sequence representation. Additionally, the bidirectionality of the architecture ensures context awareness in the past and future sequence time steps.

BERT has been demonstrated to be a versatile model, capable of pre-training on masked language modeling (Masked LM) or next sentence prediction (NSP) and subsequently fine-tuned for various tasks such as text classification, named entity recognition, and question answering. Several adaptations and similar models have emerged, including RoBERTa [63], DistilBERT [64] offering a lighter and faster alternative, and ALBERT [65] which utilizes parameter sharing to reduce model size. There are also domain-specific adaptations like ClinicalBERT [66] for medical texts, SciBERT [67] for scientific literature, LegalBERT [68] for legal documents, FinBERT [69] for financial discourse, and BioBERT [70] for biomedical text. Given BERT’s proven versatility and success across multiple disciplines and tasks, it is a fitting choice for adapting to model aptamer-target interactions in our study.

### 4.2 Preprocessing of the aptamer target pairs for BERT pre-training

The aptamer-target pairs are encoded into high-level representations suitable for the BERT model (see figure 2). Initially, the aptamer and target sequences are separately kmerized. Kmerization divides sequences into smaller non-overlapping subsequences. Due to their shorter lengths aptamers and small molecules are simply split into 1-mers (k and shift set to 1 in algorithm 1), whereas proteins are kmerized into 6-mers (k and shift set to 6 in algorithm 1). Equation 1 computes the number of kmers *θ* for a sequence given the length of the sequence *l*, the kmer size *k*, and shift *δ*:

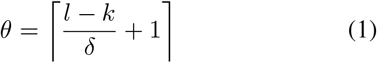

**Figure 2.**
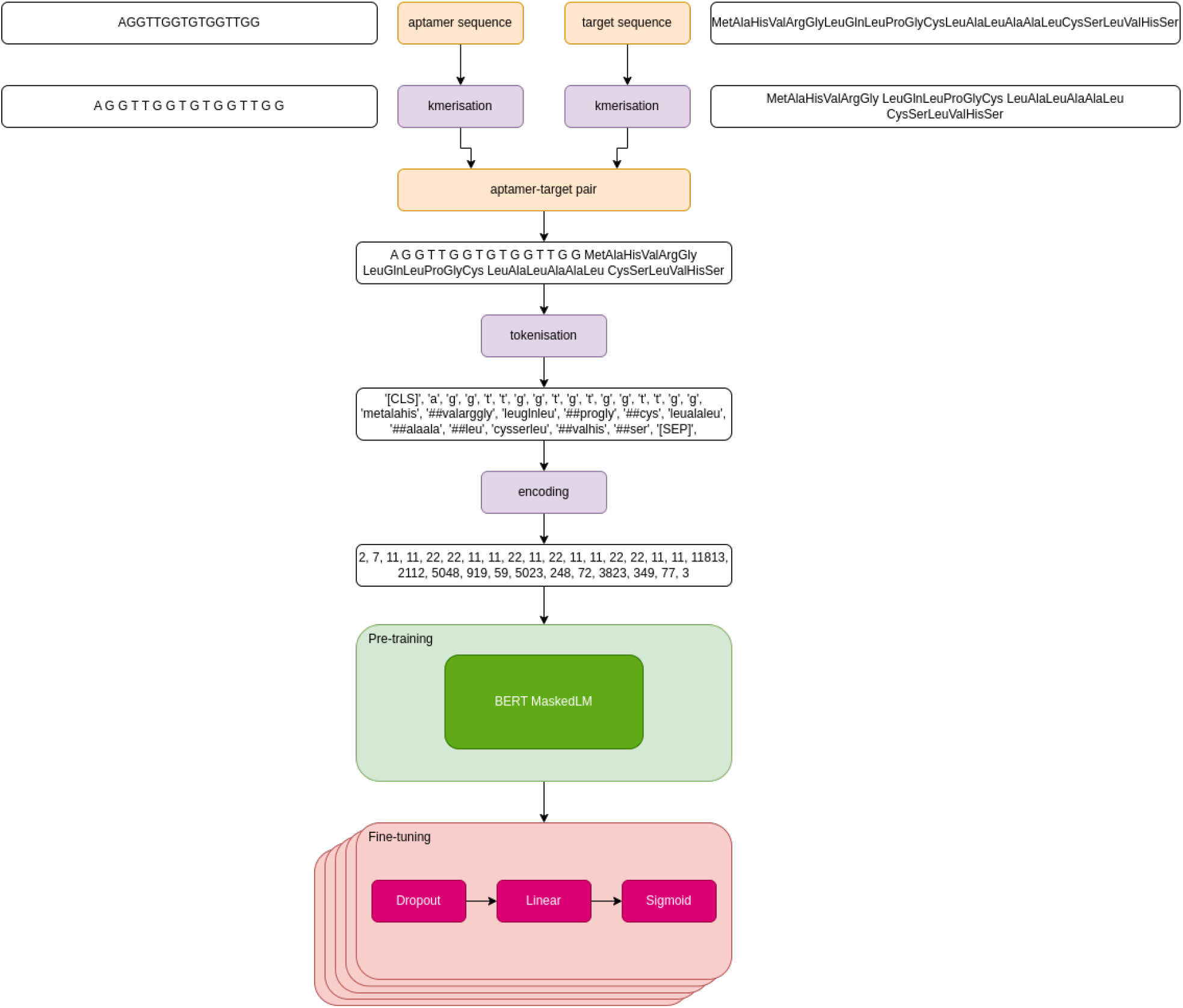
Pre-processing and model architecture

Subsequently, the WordPieceTokenizer is trained on a vocabulary containing all possible combinations of aptamers, SMILES strings and amino acids given a value of k (kmer size) from 1 to 6 to account for combinations where the last kmer of a protein contains less than 6 amino acids. We use the BERT WordPieceTokenizer [17, 30] provided by the Hugging Face library [71]. A notable advantage of the WordPieceTokenizer is its capacity to effectively handle rare “words” (in our case kmers of aptamers and targets) by breaking these kmers into subwords during training. An example of this process can be seen in figure 2. The tokenizer is trained using a vocabulary of 30000 tokens.

Since the BERT model allows for up to 512 tokens, shorter sequences are padded using the special [PAD] token, while longer sequences are truncated. After training the WordPieceTokenizer and tokenizing the input sequence, the tokens are encoded into integers. Figure 2 gives a schematic overview of the procedure of kmerization and subsequent tokenization of the aptamer and protein sequences. For small molecule targets, SMILES [72] strings are processed in the same manner using kmerization and tokenization.

### 4.3 Pre-training

During pre-training the BERT model [30] with sequence masking is trained on the complete dataset, comprising 67552 sequences as described above. The bidirectional nature of the model allows each token to interact with tokens to the left and right of its position. To prevent the current token from seeing itself during the masked language task, 15% of the tokens are randomly replaced with the [MASK] token (80%), a random token (10%), or the original token (10%). The original token is added in order to have the correct token available during the fine-tuning task since the [MASK] token only appears during the MaskedLM pre-training task. Consequently, during pre-training the sequence with masked tokens is compared to the actual sequence using the cross-entropy loss. BERT takes an input sequence of 512 tokens representing the input text. This text can be arbitrary and can represent a sentence, or, like in AptaBERT, aptamer-target pairs. Moreover, each sequence starts and ends with the special tokens [CLS] and [SEP]. The linear layers for the encoders are set to hidden dimension in 768 and the BERT model is trained for 100 epochs. The batch size is set to 16 and the dataset is split into a training set (80%) and a validation set (20%). Therefore, each batch had the dimensions 16 × 512 tokens. Data pre-processing and tokenization are conducted as described above (figure 2). For the hyperparameters, the default values as described in [30] are kept if not stated otherwise. The model is pre-trained on one Nvidia RTX 4090 GPU for approximately 36 hours.

### 4.4 Fine-tuning

After pre-training the model in a self-supervised fashion, we proceed to fine-tune it using only the labeled datasets (see figures 1a and 1b). The fine-tuning component of AptaBERT comprises a simple model containing a combination of a Dropout layer [73], followed by a Linear layer, and lastly a Sigmoid layer computing values within [0, 1] for binary classification (figure 2). We perform a 90% train 10% test split of the dataset in a random stratified manner to maintain the class balance in each subset. Subsequently, we independently train the model 5 times for 15 epochs (see figure 2, magenta boxes), with each training run taking approximately 10 minutes. Lastly, we aggregate the five individually trained models to build a deep ensemble model formulating the final classification model.

### 4.5 Ensemble learning

Both AptaBERT Prot and AptaBERT Chem consist of the same pre-trained BERT MaskedLM model combined with a deep ensemble of five fine-tuned models. These deep ensembles are built by fine-tuning five independent models, each initialized with different random seeds, on the same dataset. We consider the parameter weight vectors *w*_*i*_ of the fine-tuned models as Monte-Carlo samples of the posterior approximation, denoted by equation 2, where *w* is the weight vector of the posterior approximation *q*_E_(**w**) and *δ* is Dirac’s delta function. The predictive distribution is approximated using equation 3, where *t*^*^ are the predicted targets, *t* are the observed training targets, *x*^*^ is the data point subject to classification, and *S* is the number of independently trained models. In practice, we train *S* models − 5 in this case − with uniform weighting and take the average of the predictions to approximate the predictive distribution. The aim of the ensemble is to utilize approximate Bayesian marginalization to improve the model’s accuracy, calibration, and to reduce its variance [74, 75].

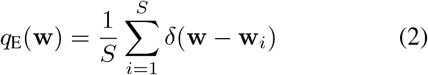

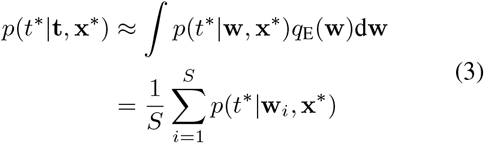

Final classification is performed by calculating the average of the estimated outputs from each of the independent models. The average estimates constitute values within the [0, 1] interval that defines the class proportions of a Bernoulli distribution. We then use the averaged predictions as the probability parameter of the Bernoulli distribution. This allows to sample a class prediction − 1000 times per data point in this study − to obtain a measure of uncertainty for the predictions.

## 5 Results

### 5.1 AptaBERT outperforms existing models

We assess the accuracy of the AptaBERT Prot model by comparing against 5 state-of-the-art models using the supervised dataset with protein targets (see Section 3). Additionally, we also report the prediction metrics for the AptaBERT Chem model using the supervised dataset with chemical targets. Note that the AptaBERT Chem model is not compared against other models due to the absence of them, therefore we only report the benchmark results using the protein target dataset. The results can be found in Table 1.

**Table 1:** Results of the AptaBERT Prot and AptaBERT Chem and benchmarking models. For the precision, recall and F1-score the weighted average is shown. The prediction decision boundary is set to 0.5. Mean and standard deviation are shown (n=5).

| Model | Accuracy | Precision | Recall | F1 score | Average Precision |
| --- | --- | --- | --- | --- | --- |
| AptaBERT Prot | $0.91 \pm 0.00$ | $0.91 \pm 0.00$ | $0.91 \pm 0.00$ | $0.91 \pm 0.00$ | $0.96 \pm 0.00$ |
| AptaBERT Chem | $0.83 \pm 0.01$ | $0.84 \pm 0.00$ | $0.83 \pm 0.01$ | $0.83 \pm 0.00$ | $0.85 \pm 0.00$ |
| AptaNet Ensemble | $0.54 \pm 0.00$ | $0.30 \pm 0.00$ | $0.54 \pm 0.00$ | $0.38 \pm 0.00$ | $0.84 \pm 0.01$ |
| AptaNet [26] | $0.70 \pm 0.03$ | $0.70 \pm 0.03$ | $0.70 \pm 0.03$ | $0.70 \pm 0.03$ | $0.82 \pm 0.01$ |
| PPAI [44] | $0.77 \pm 0.03$ | $0.77 \pm 0.03$ | $0.77 \pm 0.03$ | $0.77 \pm 0.03$ | $0.81 \pm 0.03$ |
| Li et al. [45] | $0.85 \pm 0.03$ | $0.85 \pm 0.03$ | $0.85 \pm 0.03$ | $0.85 \pm 0.03$ | $0.81 \pm 0.03$ |
| Zhang et al. [47] | $0.70 \pm 0.04$ | $0.70 \pm 0.04$ | $0.70 \pm 0.04$ | $0.70 \pm 0.04$ | $0.71 \pm 0.03$ |

Benchmarking is performed by following the model settings and data preprocessing described by the authors. We also use the same train-test split for all the models and run 5 instances of each model initialized with different seeds. The random forest based methods rely on a combination of complex sequence feature extraction methods including pseudo-amino acid and pseudo-nucleic acid composition as well as other amino acid and nucleic acid physicochemical properties (see section 2.1) that constitute their input to the model. On the other hand, recall that AptaBERT Prot uses aptamer and protein sequence kmerization with subsequent tokenization (see Section 4.2) that is funneled into an embedding representation that summarizes the interacting pair of sequences.

The AptaBERT Prot model achieves the best scores for all the analyzed metrics, with 91% accuracy, precision, recall

### 5.3 AptaBERT Prot predicts interactions with high certainty

Figure 3 shows the predictions of the three ensemble models AptaBERT Prot, AptaBERT Chem and AptaNet Ensemble. The x-axis shows the indices of the samples in the test set sorted in ascending order by the predicted value *t*^*^ ∈ [0, 1] and the y-axis shows the probability scores. Orange points denote the true target values 0 (negative and F1-score as well as an average precision score (APP) of 96%. In comparison, the random forest-based models Li et al. [45], Zhang et al. [47] and PPAI [44] attained lower scores for the same metrics with Li’s model presenting the second best scores among the lineup.

**Figure 3.**
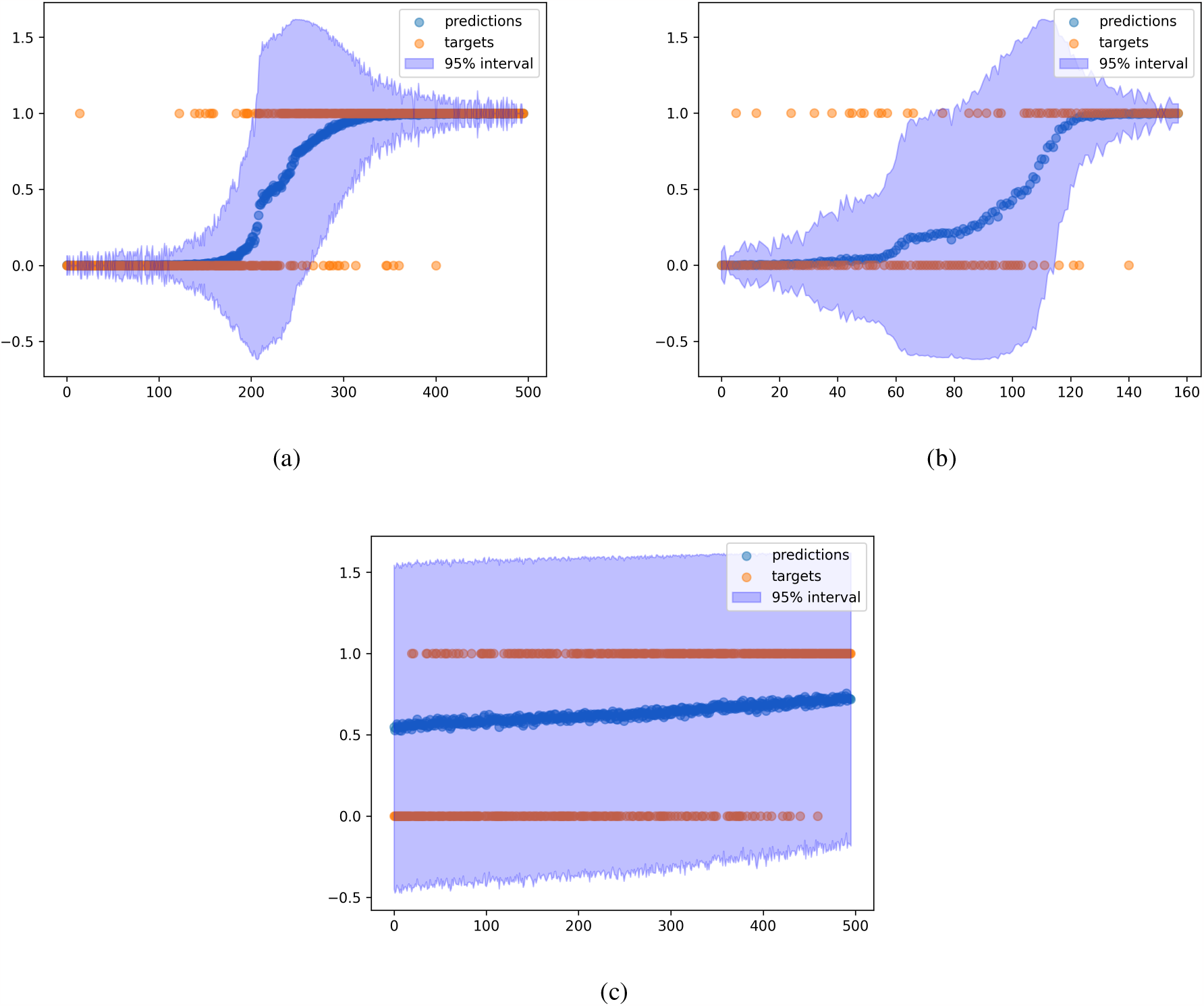
Predictive distributions of a) AptaBERT Prot, b) AptaBERT Chem and c) AptaNet. The x-axis denotes the indices of the test samples sorted by ascending prediction value. The y-axis denotes the values. Orange points present the true labels, blue points denote the average prediction scores drawn from the predictive distribution and the blue area denotes the 95% confidence interval of each prediction.

The results show how classical ensemble models, such as the random forest, sustain a large classification power while evading the large computational training times from the deep learning ensemble. AptaBERT Prot requires a pretraining time of 36 hours and 10 minutes per fine-tuning model (15 epochs), whereas random forests can be trained within 1 minute. Nonetheless, AptaBERT displays superior performance compared to the benchmarked models indicating a potential to capture the biochemical interactions more accurately.

### 5.2 Comparison against existing deep learning architectures

To assess the benefit of Large Language Models over simpler deep learning architectures, we compare AptaBERT against AptaNet, the only available neural network based model, to our knowledge, trained for the aptamer-protein interaction task. AptaNet [26] is a multi-layer perceptron (MLP) that also uses similar physicochemical sequence encodings to those utilized by the the random forests. These features subsequently undergo selection using a random forest feature selector.

First, we run AptaNET 5 times independently with different random seeds achieving an average precision (AP) score of 0.85, while other metrics show scores around 0.7 (refer to table 1). Notably, AptaNet consistently made accurate predictions for samples belonging to the positive class, whereas its accuracy drops significantly for the negative class.

To ensure a balanced comparison with our AptaBERT Prot model, we proceed to train AptaNet as an ensemble model (AptaNet Ensemble). We observe that AptaNet Ensemble shows poor accuracy, precision, recall, and F1-scores at the 0.5 decision threshold whereas the class-balance insensitive metrics Average Precision and ROC-AUC are higher. These results suggest that the model is more accurate at a different threshold and its performance drops when approaching negative class designation. We further analyze this hypothesis in section 5.3 class, no aptamer target interaction) and 1 (positive class, aptamer target interaction), while blue data points denote the predictions according to equation 3.

We proceed to calculate the 95% confidence interval of the Bernouilli distribution by using the Wald interval, an approximation using the Normal distribution [76]. As discussed in a previous section, we are interested in comparing our deep learning model with another deep learning architecture AptaNet. Figures 3a and 3c show the results of the two ensembles quantifying the uncertainty of the predictions. AptaBERT Prot is certain about predictions for samples with indices between 0-150 (negative class) and 300-500 (positive class), while the uncertainty is larger for samples between 150-300.

These results indicate that the five independent models predict these data points to be in different classes, contributing to uncertain predictions with a spread between approximately -0.5 and 1.5 for samples with indices around 200 (figure 3a). On the other hand, for data points with narrow confidence intervals, the different models predict values close to each other and thus agree on the predicted label more often.

### 5.4 AptaBERT performs on protein and small molecule targets

BERT models are very versatile due to their self-supervised pre-training capabilities. Thus, we pre-trained AptaBERT on both aptamer-protein, and aptamer-small molecule pairs simultaneously (figure 1c), albeit with only 7.34% samples containing small molecules. Therefore, we investigate the performance of the AptaBERT Chem model which is fine-tuned on a labeled dataset containing only aptamer-small molecule pairs. Despite the much smaller pre-training and fine-tuning datasets, AptaBERT Chem achieves scores of 83% and higher for all metrics (figure 1) on a test set with a label balance of 43% positive to 57% negative samples (1b). Like for the other two deep learning models, we also investigate the uncertainty of this model’s predictions in figure 3b. Among the 160 samples in the test set for AptaBERT Chem, the uncertainty of most of the samples is acceptable and did not reach the threshold of 0.5 for either true label. However, some samples with indices between 60 to 110 show high uncertainty, indicating disagreement between the five individual models. Bigger datasets during pre-training and fine-tuning will likely improve the performance of both AptaBERT models.

### 5.5 Sequence embeddings showcase better performance than feature-based methods

The results discussed above highlight the advantage of pretraining on a larger dataset to learn general patterns within the sequences compared to solely training on extracted features of a smaller dataset (table 1). AptaBERT uses simple embeddings based on kmerization and subsequent tokenization of these kmers in combination with BERT, a large 110M parameter transformer model learning patterns from “words” in a “sentence”. This stands in stark contrast with the methods by other groups (table 1) which designed intricate algorithms to craft and extract features from frequencies and physicochemical properties of the sequences in combination with smaller models (Random-Forests and MLPs). As we can see in figure 4a, the overall performance of our AptaBERT Prot model is undisputed, scoring a receiver operator characteristic area under the curve (ROC-AUC) of 0.96 compared to scores of 0.81 and lower for the benchmarking models. The ROC-AUC is the area under the curve when plotting the false positive rate (precision, x-axis) against the true positive rate (recall, y-axis) at different thresholds of the decision boundary of the binary classification and it measures the model’s ability to discriminate between the positive and negative class, where a ROC-AUC of 1 means a perfect model and 0.5 indicates a random choice.

**Figure 4.**
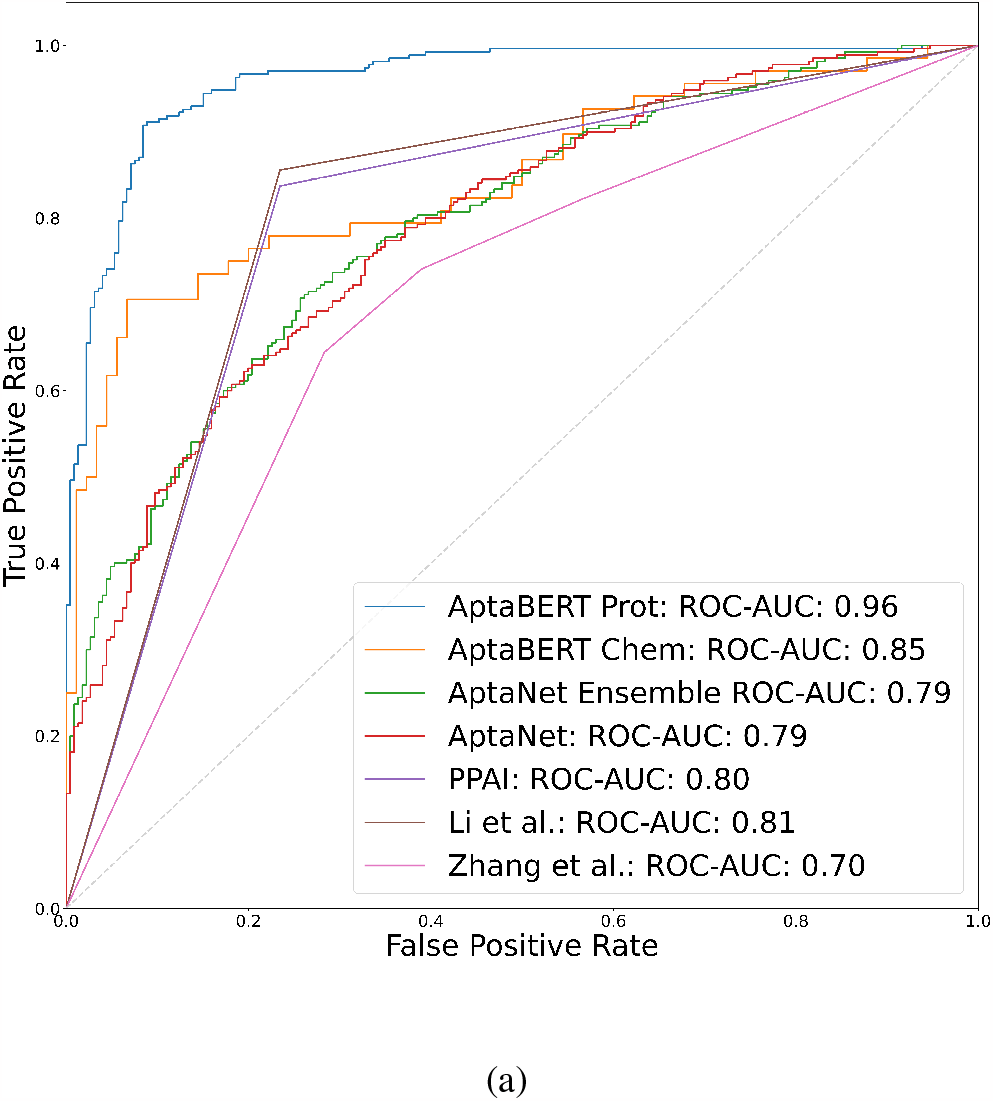
Receiver Operator Characteristic Area Under the Curve (ROC-AUC) of the different models.

These findings suggest that AptaBERT Prot is a superior, albeit much more resource-consuming, model. The combination of kmer encodings of aptamer-target pairs together with tokenization and BERT masked language modeling based on the transformer architecture [30, 55] improves the model’s ability to distinguish between the positive and negative class. Nevertheless, especially, the model by Li et al. [45] showed good performance as well, given a ROC-AUC of 0.81 with their random forest based model.

### 5.6 Attention representations reveal aptamer-target interactions

To assess if the model’s predictions are biased towards only learning from either the aptamer or the protein features and not the aptamer-target interactions, we analyze the attention layer’s outcomes. Alongside, we compare them against the feature importance of the best-performing random forest model by Li et al. [45]. In transformer models such as AptaBERT Prot, we can investigate the attention weights of the different layers, and in particular the last layer, to unravel the importance of aptamer and protein interactions. In figure 5 we plot the attention weight heatmaps of two aptamer-protein pairs (figures 5a and 5c) as well as the average attention per matrix quadrant (figures 5b and 5d). Brighter colors (more yellow) indicate higher values and thus higher attention values. There are four quadrants separated by dashed red lines which separate the areas of aptamer-aptamer (left upper), aptamer-protein (right upper), protein-aptamer (left lower) and protein-protein (right lower) attentions. Notice that there are two quadrants of aptamer-protein interaction and one each for the feature importance of aptamers and proteins. These attention heatmaps are obtained by averaging the 12 self-attention heads of the 12^*th*^ attention layer, minmax scaling and taking the 95% quantiles (two-sided) of the data. To obtain a quantifiable representation of the attention weights, we compute the average of each of the the aptamer-aptamer (*Aptamer* column), protein-protein (*Protein* column) and the two aptamer-protein quadrants (*Interaction* column) as a fraction of the total of all mean values (all quadrants) (figures 5b and 5d). We can see in figures 5a and 5c that the token lengths of aptamer and protein are 100 and 118 tokens and 22 and 54 tokens respectively. In both of these heatmaps, the quadrants with intra-molecular interactions show high attention values, while the lower left and upper right quadrants, representing the interaction between the two molecules, have lower attention values. In the shown example, the attention weights over the aptamer representation corresponds to 41% of the mean attention values which in this case equals the attention given to the protein. The remaining 18% account for the inter-molecular interaction between the aptamer and the protein (figure 5b). In figure 5d, to assess possible length biases, we select an example where the size ratio between aptamer and protein tokens differs (22 aptamer tokens vs. 54 protein tokens). We show that in this case, the aptamer importance appears higher than the protein’s, with values of 45% and 35% respectively, whilst maintaining awareness of the inter-molecular interactions, which account for 20% of the average attention. Overall, we present a quantifiable method that can help assess the reliability of the model predictions by estimating a feature importance measurement based on the attention weights.

**Figure 5.**
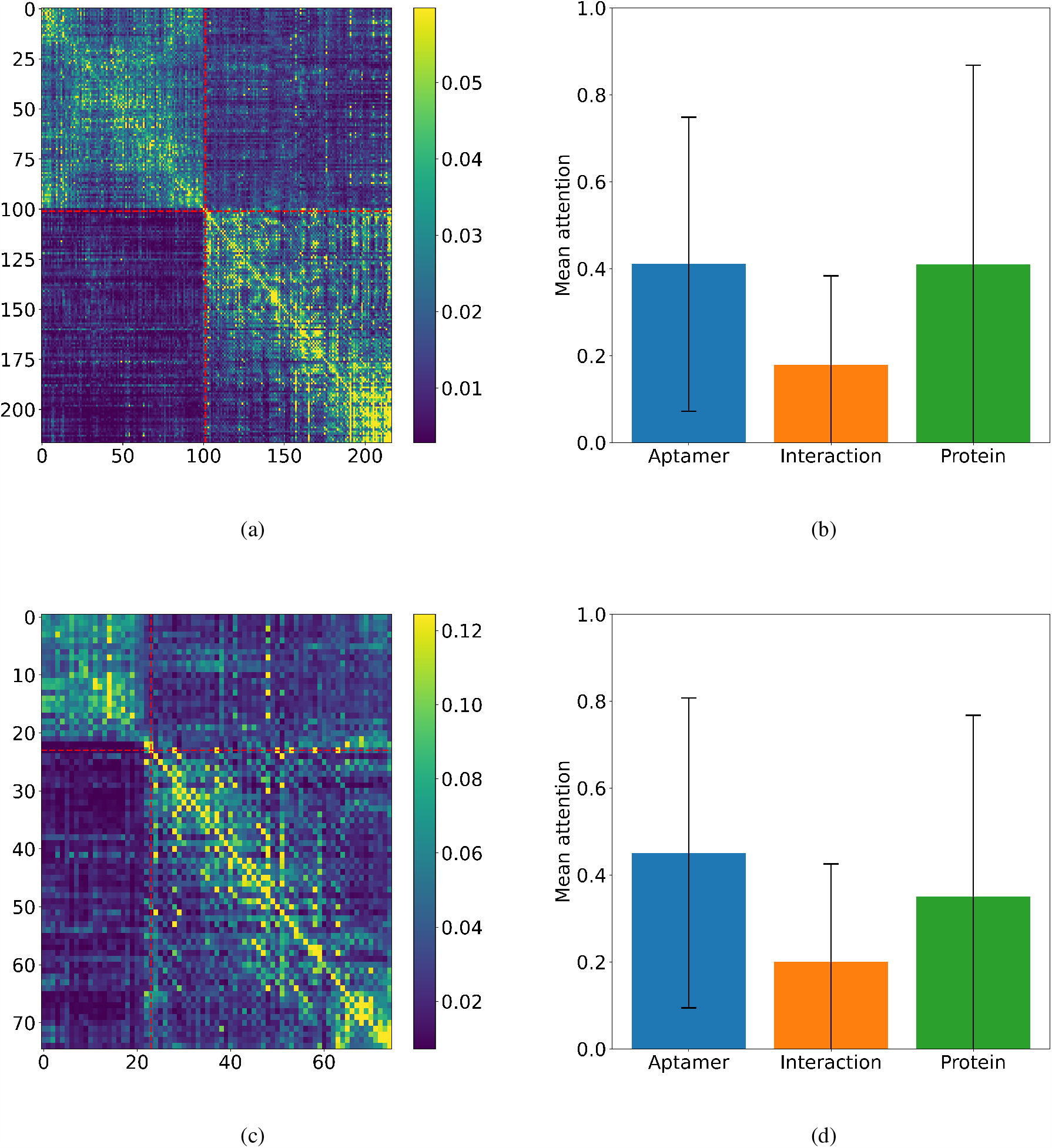
Average attention of AptaBERT Prot of two aptamer-protein pairs. a) and c) heatmap of the 12^*th*^ attention layer. The 12 self-attention heads of the 12^*th*^ attention layer are averaged, min-max scaled and the 95% quantiles are used. The red dashed lines show the four different quadrants. Upper left: aptamer-aptamer attention. Upper right: aptamer-protein attention. Lower left: protein-aptamer attention. Lower right: protein-protein interaction. The x-axis and y-axis show the indices of the tokens of the aptamer-protein pairs (special tokens are omitted). b) and d) Mean attention of the quadrants as a fraction of all quadrants’ attention weights. The mean attentions of the aptamer-protein and protein-aptamer quadrants are shown together as the *Interaction* column, while the mean aptamer-aptamer attentions and mean protein-protein attentions are named *Aptamer* and *Protein* respectively. Mean and standard deviation are shown.

Next, we investigate whether AptaBERT captures long range sequence interactions thanks to the presence of fully connected layers and positional encodings that can account for them. This is a key difference between deep learning models and other machine learning classifiers such as random forests. Random forests treat each position in the input (i.e. sequence or feature vector) as independent, failing to capture any possible interaction between distant positions [77]. This entails that their predictions are solely based on a summary representation of each of the sites of the sequences independent from each other. To show how, AptaBERT takes into consideration aptamer-target interactions, we compute the average attention for the entire *Protein* dataset (figure 6a) and compare it to the feature importance results of the random forest model by [45] (figure 6b). We find that the average attention for the aptamers, aptamer-target interactions and proteins of AptaBERT Prot are 46.5 ± 6.1%, 20.8 ± 2.2%, and 32.7 ± 5.1% respectively. On the other hand, while our model obtains relatively balanced results between the different features, the model by Li et al. puts 89% of its emphasis on proteins and solely 11% on the aptamers. The random forest model cannot account for aptamer-target interactions and neither uses an explicit representation for the interaction between aptamers and proteins. Figure 6c shows the feature importance in decreasing order of all 220 features of the random forest with protein features highlighted in orange and aptamer features in blue. The first 16 features represent protein properties and account for 36% of the overall feature importance. The aptamer characteristics seem to account solely for 11% and are additionally not among the more important features (figure 6c, blue features). The great emphasis on the protein features can also be explained with the fact that the model mostly used protein features and fewer aptamer features [45]. Nevertheless, in comparison to AptaBERT, we can see in figure 6c that the predictive power of the random forest model is mostly explained by protein features, while aptamer properties are less important. Additionally, no inter-molecular interaction insights can be gained from the random forest model.

**Figure 6.**
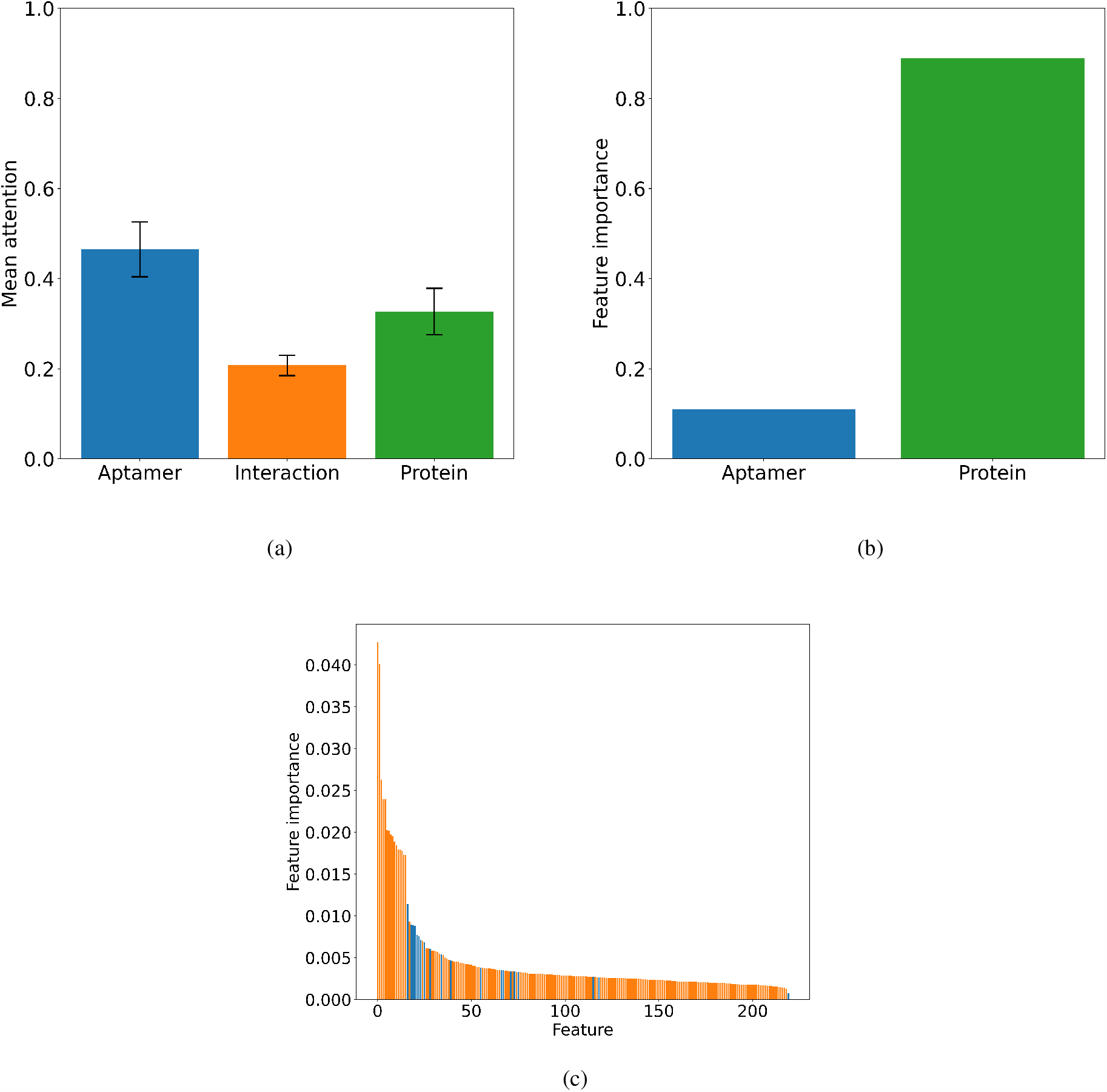
Comparison of AptaBERT Prot mean attention and the feature importance of the random forest model by Li et al. [45]. a) shows the mean attention of the quadrants as a fraction of all quadrants’ attention weights for all samples of the *Protein* dataset with standard deviation. The 12 self-attention heads of the 12^*th*^ attention layer are averaged and min-max scaled and the 95% quantiles are used. b) shows the feature importance of the aptamer and protein property contributions for the random forest model [45]. c) shows the feature importance of each feature of the random forest model.

## 6 Conclusion

The ability to accurately predict aptamer-target interactions holds immense potential for biomedical applications in both diagnostics and therapeutics. To further the research in this field, we present AptaBERT, a BERT-based deep learning model. It is pre-trained using self-supervised learning on unlabeled aptamer-target pairs. We then fine-tune this model as deep ensemble on two classification tasks: Predicting aptamer-protein and aptamer-small molecule interactions. AptaBERT achieves superior performance with a ROC-AUC of 96% for AptaBERT Prot, outperforming all benchmarking models by at least 15%. Additionally, we successfully demonstrate a method to assess the feature importance of AptaBERT by quantifying the interactions between aptamers and proteins using AptaBERT Prot’s attention layers and showcase that our model learns from both, aptamers, proteins and their interactions. Moreover, to our knowledge, AptaBERT is trained on the largest aptamer-target interaction dataset, containing 67552 interaction pairs and we anticipate to refine the model’s results with Dianox’s growing dataset.

Besides offering more accurate predictions, this large dataset also enables the potential of extending the architecture to a generative model, assisting in molecular design. This could allow for the generation of aptamers against many yet untargeted proteins. Using similar deep learning architectures, it is possible to design novel aptamer molecules for specific target molecules in silico, presenting a time- and resource-efficient alternative to traditional laboratory-based methods like SELEX. We aim to further improve the model to incorporate interactions in the 3D molecular space, providing deeper insights into the molecular mechanisms behind aptamer binding. This advancement may enable the identification of small molecules effective against druggable RNA structures. With this study, we show that the combination of a large curated dataset and modern deep learning models yields accurate aptamer-target interaction predictions.

## Appendices

### A Equations

#### A.1 Sigmoid function

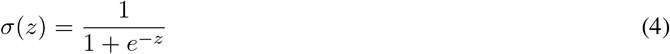

#### A.2 Classification metrics

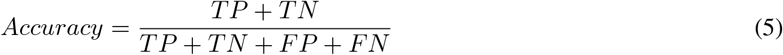

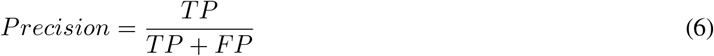

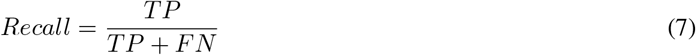

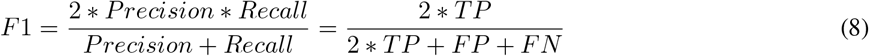

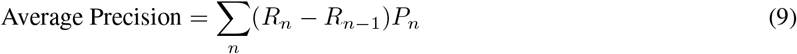

#### A.3 Binary cross-entropy

**Variables:**

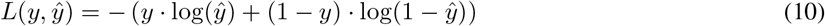

- *L*(*y, ŷ*): Binary cross-entropy loss.
- *y*: True binary label (either 0 or 1).
- *ŷ*: Predicted probability of the sample belonging to class 1 (the range is [0, 1]).

#### A.4 Min-Max Scaling

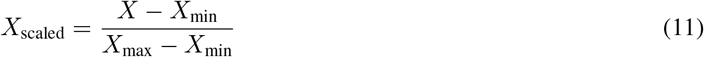

### B BERT hyperparameters

**Table 2:** Common BERT Hyperparameters and Default Values.

| Hyperparameter | Description | Default Value |
| --- | --- | --- |
| Vocabulary Size | Size of the vocabulary | 30000 |
| Hidden Layers | Number of transformer layers | 12 |
| Hidden Units | Number of hidden units in each layer | 768 |
| Attention Heads | Number of attention heads in each layer | 12 |
| Max Sequence Length | Maximum input sequence length | 512 |
| Batch Size | Number of training examples in one batch | 16 |
| Learning Rate | Step size during optimization | 5e-5 |
| Dropout Probability | Probability of dropout | 0.1 |
| Warmup Steps | Number of steps during learning rate warm-up | 10% of total training steps |
| Weight Decay | L2 regularization strength | 0.01 |
| Optimizer | Optimization algorithm | Adam |
| Maximum Position Embedding | Maximum position index for embeddings | 512 |
| Initializer Range | Range of values for weight initialization | 0.02 |
| Gradient Accumulation Steps | Number of steps before gradient update | 1 |

### C Kmerization algorithm

#### Algorithm 1

Kmerization algorithm for sequences

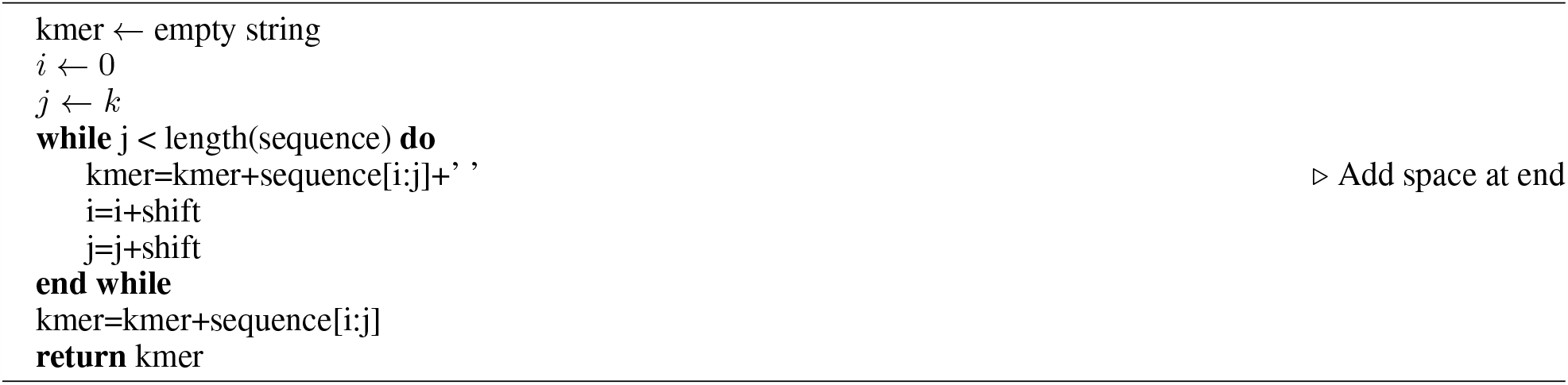

### D AptaBERT loss curve

**Figure 7.**
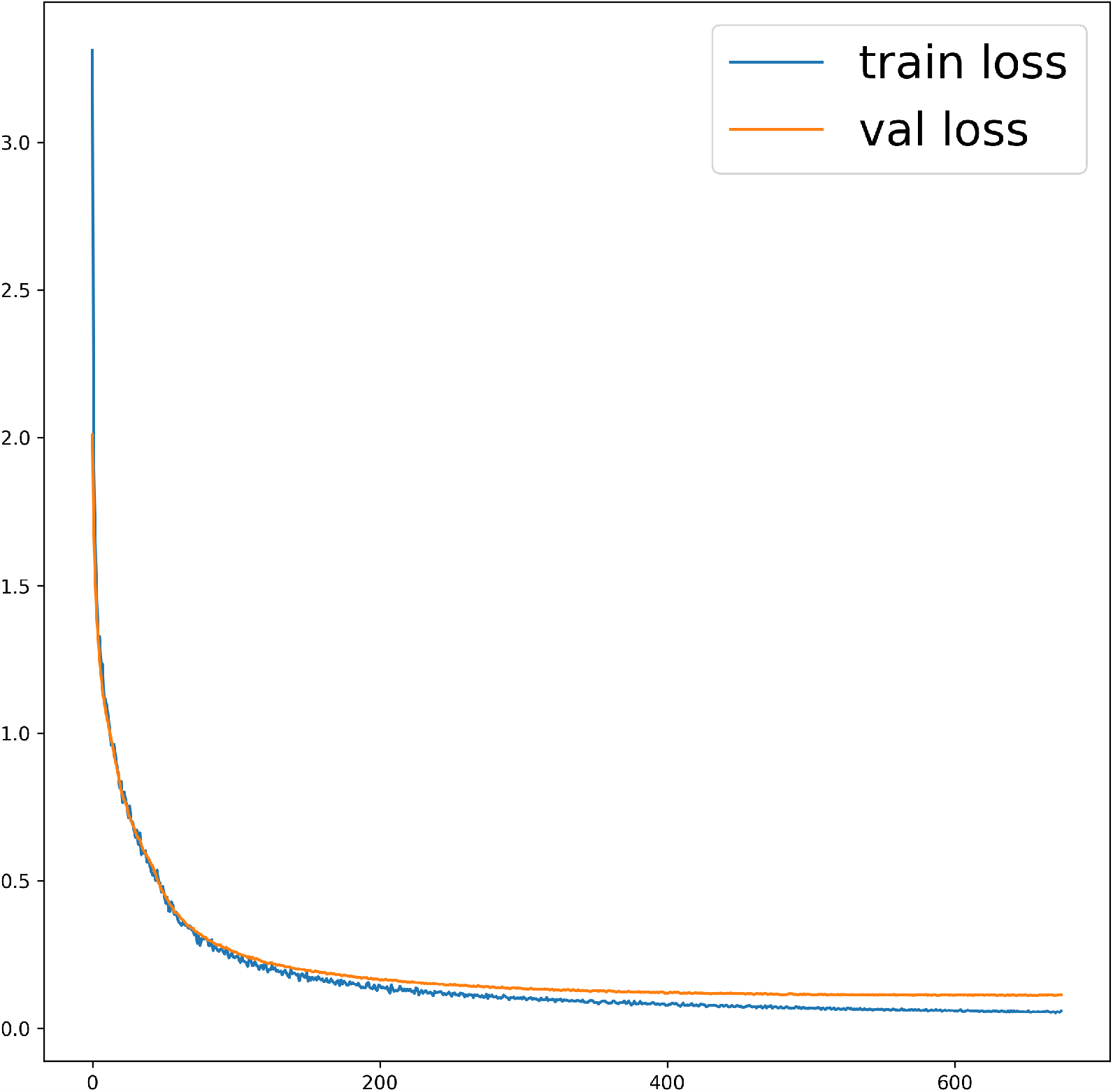
AptaBERT loss curve. The model was trained for 100 epochs or 337500 steps. Among these steps 675 data points with losses were recorded.

